# Light pollution affects the behavior and life history traits of aquatic invertebrates

**DOI:** 10.1101/2025.10.09.679630

**Authors:** Enav Bereza, Ida Rotter, Sofia Paraskevopoulou, Frida Ben-Ami

## Abstract

Circadian rhythms regulate essential biological processes, including behavior, reproduction and survival, across diverse organisms. Disruptions to this cycle, whether through artificial light at night (ALAN) or reduced exposure to full-spectrum daylight, can interfere with biological processes and ultimately threaten biodiversity. This concern is growing as ALAN spreads beyond urban areas through streetlights and skyglow. Aquatic ecosystems, in which zooplankton play a key role as primary consumers as well as prey for higher trophic levels, are particularly vulnerable. Despite growing research on the impact of ALAN on some zooplankton, rotifers remain largely understudied. Here, we conducted two experiments to assess the effects of light on *Brachionus* species. The first experiment examined how different ALAN wavelengths influenced life history traits in three *Brachionus* species, while the second one investigated the attachment behavior of *B. rubens* on a *Daphnia* host under four light conditions. Our findings reveal that ALAN has strong but species-specific effects on rotifers. Green light enhanced survival and reproductive output in *B. calyciflorus* sensu stricto, while suppressed reproduction in *B. fernandoi*. In *B. rubens*, white light particularly altered reproductive patterns and attachment behavior. These results highlight the complex and species-specific impacts that ALAN has on rotifers, emphasizing the need for further research to fully understand its broader ecological consequences.

## Introduction

Light is one of the most fundamental environmental cues shaping life on Earth. It regulates primary production, circadian rhythms and seasonal cycles, thereby influencing physiology and behavior across taxa (Chen et al., 2020; Stewart & Albrecht, 2025; Wahl et al., 2019; Wu et al., 2025). Yet, the natural light-dark cycle is increasingly being disrupted by artificial light at night (ALAN), also known as light pollution, which affects both flora and fauna with profound ecological consequences (Bennie et al., 2016; Falcón et al., 2020; Marangoni et al., 2022; Seymoure et al., 2023). One of the primary concerns regarding light pollution is the impact on animal biodiversity, as it alters otherwise regulated processes, such as circadian rhythms and consequently disrupts behavior across taxa (Alaasam et al., 2021; Lei et al., 2024; Levy et al., 2022). Such disruptions cascade to other ecosystem processes, as for instance parasitism, with consequences for ecosystem functioning (Bolliger et al., 2020; Brown et al., 2023; Giavi et al., 2020; Knop et al., 2017; Sanders et al., 2018; Shivanna, 2022).

While the effects of light pollution on vertebrates, birds and insects have been well documented (Alaasam et al., 2021; Bennie et al., 2016; Falcón et al., 2020; Lei et al., 2024; Levy et al., 2022; Marangoni et al., 2022; Seymoure et al., 2023), the impact on microscopic aquatic organisms, particularly zooplankton, remains largely unexplored and is mostly confined to aquaculture species that serve as food for larval fish. Experimental work has shown that light pollution can interfere with key zooplankton traits. For instance, exposure to different wavelengths affected the development, reproduction and antipredator defenses of *Daphnia magna* (D. Li et al., 2022). Additionally, *D. longispina* exhibited clone specific responses to light pollution depending on the spectral composition of the light source, with more adaptive depth selection occurring under LED and high-pressure sodium light (Maszczyk et al., 2021). Given their crucial ecological role as primary consumers, prey for higher trophic levels, and contributors to nutrient cycling (Gomes et al., 2019; Sterner, 2009; Thorpe, 2020), disruptions in zooplankton behavior and life history may have cascading consequences for aquatic ecosystems.

In this study we focus on rotifers, a key but understudied zooplankton group in light pollution research. Light shapes several life history and behavioral traits in rotifers, including dormancy, hatching and reproduction (Dupuis & Hann, 2009). For instance, light intensity and wavelength can alter the timing of hatching (Dupuis & Hann, 2009), and affect reproductive success (e.g., phototactic responses to UV light differ between sexes in the *Brachionus calyciflorus* complex; Colangeli et al., 2019). Despite the profound effects of light, most studies of light pollution in rotifers have focused on species used in aquaculture, such as *Brachionus plicatilis* and *B. manjavacas* (Kim et al., 2013; Kim, Sawada, Hagiwara, 2014; Wang et al., 2024). These studies show that light wavelength and intensity influence phototaxis, reproduction and resting egg production (Kim et al., 2013; Kim, Sawada, Hagiwara, 2014). Moreover, light pollution can interact with other factors, such as food availability, as shown by contrasting effects of purple and white ALAN on *B. plicatilis* development depending on diet (Wang et al., 2024). These findings suggest that light pollution could strongly shape rotifer ecology, yet its effects remain poorly understood outside of aquaculture relevant species.

The *B. calyciflorus* species complex consists of four closely related taxa (Michaloudi et al., 2018; Papakostas et al., 2016), which have evolved different physiological responses to environmental stressors such as temperature (Paraskevopoulou et al., 2018, Paraskevopoulou et al., 2020). Since temperature and photoperiod are linked, these species may also differ in their sensitivity to light. Given that *B. calyciflorus* sensu stricto (s.s.) is typically considered a summer species (Y. Zhang et al., 2018), it may have adapted to natural light fluctuations, making it potentially less sensitive to light pollution. In contrast, *B. fernandoi*, which is considered a winter species, might be adapted to lower-light conditions and thus, may be more vulnerable to light pollution. Based on this, we hypothesized that light pollution may have stronger effects in some *B. calyciflorus* species than in others. Another widespread rotifer, *B. rubens*, attaches to surfaces or the exoskeleton of other organisms, particularly cladocerans, which enhances feeding and predator avoidance (Gilbert, 2019; Green, 1974; Iyer & Rao, 1995; May, 1989). This attachment behavior varies depending on endoparasite infections, highlighting the complexity of biotic–biotic interactions (Hirshberg & Ben-Ami, 2025). Light pollution may disrupt this attachment behavior by altering their phototactic responses. If *B. rubens* detaches more frequently due to altered light cues, it may experience reduced feeding efficiency or increased predation risk. These disruptions can potentially affect population dynamics and trophic interactions within aquatic ecosystems, altering the stability of planktonic communities.

Here, we tested how ALAN influences life history traits and behavior in three freshwater rotifers: *B. calyciflorus* s.s., *B. fernandoi* and *B. rubens*. We hypothesized that the effects of ALAN would differ among species, reflecting their ecological adaptations, and that altered light cues could disrupt the attachment behavior of *B. rubens*. By addressing these questions, we aim to better understand how light pollution affects rotifers beyond aquaculture species, and to explore potential cascading consequences for freshwater ecosystems.

## Materials and methods

### Experiment 1: Effects of light pollution on life history traits of rotifers

Dormant eggs of *B. fernandoi* and *B. rubens* were collected during a survey of freshwater ponds in Israel (Hirshberg et al., 2023), while the *B. calyciflorus* sensu stricto clone originated from Egelsee, Austria. All species were maintained under standard laboratory conditions (12:12 light:dark cycle at 21°C). Cultures were reared in jars containing 300 mL of artificial *Daphnia* medium (ADaM, Ebert et al., 1998; Klüttgen et al., 1994) and fed daily with the algae *Scenedesmus* sp. *ad libitum*.

To assess the effects of light pollution on rotifer life history traits, five lighting regimes were established during the rotifer’s night cycle (12h): (1) Natural photoperiod/control which received complete darkness, (2) Red light (580-740 nm, peak at 620 nm), (3) Blue light (430-500 nm, peak at 450 nm), (4) green light (460-580 nm, peak at 540 nm), and (5) broad-spectrum white light (460-700 nm, peak at 460 nm, and broader distribution between 480-640 nm; *See Supplementary material for the full spectrophotometer output*).

Females bearing subitaneous asexual eggs were isolated from the original culture, placed individually in a microtiter well and inspected thoroughly for hatched neonates. Upon hatching the neonates, representing the experimental animals, were transferred into a new well containing 1 mL algal suspension composed of *Scenedesmus* sp. (75 × 10^4^ cells/mL). For each light treatment and species, 30 individuals were recorded. All experimental groups, comprising the four light pollution and the control, were subjected to identical daytime lighting (broad-spectrum white daylight at 440-660 nm, peaks at 440, 500, 540, 580 and 620 nm) for 12 h. The only variation occurred at night: while the control group experienced darkness, each ALAN treatment was exposed to its designated nighttime light color. Survival and reproduction output were recorded every 24 h (Figure 1a). Every day, the experimental individuals were transferred into a new well with fresh food suspension. The life history experiment continued until all individuals died.

**Figure 1.**
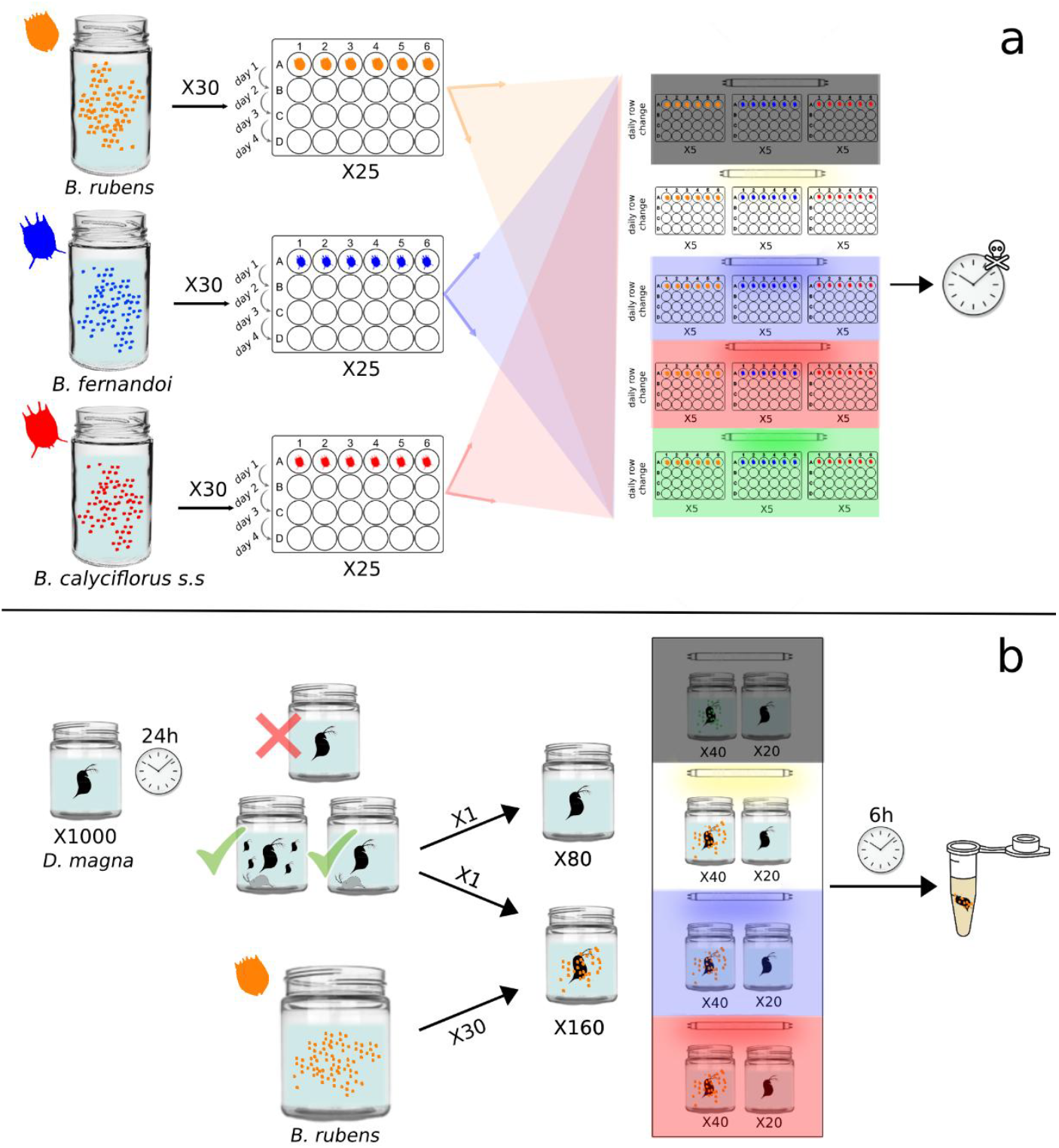
Experimental design for assessing the impact of light pollution on (a) life history traits of three rotifer species, and (b) the attachment behavior of *B. rubens*.

### Experiment 2: Effects of light on the attachment behavior of B. rubens

A cohort of *Daphnia magna* individuals (clone M10, Israel; Izhar & Ben-Ami, 2015; Izhar et al., 2015) was acclimated for four generations under a 12:12 light-dark cycle at 21°C. Experimental animals were isolated from the 3^rd^ brood of the 4^th^ acclimation generation. *D. magna* is a 2-3 mm crustacean used by *B. rubens* as a substratum for attachment (Goren & Ben-Ami, 2013; Green 1974). The *B. rubens* clone (Tel Yitzhak, Israel) was also acclimated for several generations under the same conditions. To select individuals that exhibit attachment behavior, we introduced *D. magna* to the acclimated *B. rubens* clone, removed the *Daphnia* with attached *B. rubens* and used these individuals to establish the experimental clone.

The experiment consisted of thirty *B. rubens* individuals, manually counted under a dissecting microscope (Leica M205C, Germany), and one *D. magna* individual being introduced into a jar containing 70 mL of ADaM (Ebert et al., 1998; Klüttgen et al., 1994). To avoid molting during the experiment, which could bias the attachment behavior, we selected *Daphnia* individuals that had molted and spawned recently. In total there were 160 jars containing 30 *B. rubens* and one *Daphnia*, and 100 jars containing only *Daphnia* as a control (to monitor unexpected treatment responses). During daytime, jars were exposed to four light treatments for six hours as follows: (1) daylight (440-710 nm, i.e., control), (2) dark, (3) red light (625 nm), and (4) blue light (455 nm) (Figure 1b). After six hours of exposure, *Daphnia* with attached rotifers were fixed with 6 μL of Lugol solution and rotifers were counted. At the end of the experiment, each jar was inspected for molts and offspring. *Daphnia* that molted, spawned or died during the six hours of exposure were excluded from further analysis.

### Statistical analyses

All statistical analyses were conducted using R v.4.3.1. Homogeneity of variance was assessed using Levene’s test (‘car’ package v.3.1.2; Fox et al., 2023), and residual normality was evaluated with the ‘DHARMa’ package v.0.4.7 (Hartig, 2025). When normality assumptions were violated, generalized linear models (GLMs) were applied instead of analysis of variance (ANOVA).

### Experiment 1: Effects of light pollution on life history traits of rotifers

In the life history experiment data were analyzed separately per species, as one clonal line per species was used. During the experiment we observed male-producing individuals, primarily in *B. rubens* and *B. calyciflorus* s.s. Consequently, the data were split into two datasets, the asexual (amictic) fraction of the population containing data from asexual females, and the sexual (mictic) fraction of the population containing data from sexual females. Females were characterized as sexual when they produced male offspring. In *B. fernandoi*, only a few individuals were sexual females and thus they were excluded from any further analyses. We assessed the effect of light pollution on four phenotypic traits. In asexually reproduced females we measured survival, total number of offspring as a proxy for fecundity, and fecundity per day, representing the number of offspring per day of survival. In sexually reproduced females we measured survival and the number of broods with male offspring.

For the species *B. rubens* and *B. calyciflorus* s.s. survival was analyzed using a gamma-distributed GLM, with light pollution and rotifer type (sexual or asexual) as explanatory variables. For model selection, we used the ‘*dredge’* function (‘MuMIn’ package v1.48.4; Bartoń, 2024), which selects the most parsimonious model based on cAIC. Statistical significance for our explanatory variables in the best fitting model was obtained using a Wald Chi-Square test. For the species *B. fernandoi*, which produced only a handful of sexually reproducing females, light pollution was the only explanatory variable included in the model. This model was compared to the null model using a Chi-squared test. For all three rotifer species, fecundity and fecundity per day were assessed using a GLM with a gamma distribution and light pollution as an explanatory variable. Post hoc comparisons were performed using the ‘emmeans’ package v.1.8.8 (Lenth, 2023).

In the case of *B. rubens* and *B. calyciflorus s*.*s*. that produced sexually reproducing females, we also assessed the effect of light pollution on male brood production. The number of male brood production was analyzed using a GLM with a gamma distribution and light pollution as an explanatory variable. Post hoc comparisons were performed using the ‘emmeans’ package v.1.8.8 (Lenth, 2023).

### Experiment 2: Effects of light on the attachment behavior of B. rubens

In the attachment behavior experiment, analysis of variance (ANOVA) was used to assess the effect of light pollution on the number of attached rotifers on the *Daphnia* individual. Pairwise t-tests with Tukey’s HSD correction were used for post hoc comparisons (‘stats’ package v.4.3.1).

## Results

### Experiment 1: Effects of light pollution on life history traits of rotifer species

In the life history experiment, we compared how different spectra of ALAN affected traits such as fecundity, survival and fecundity per day (total offspring divided by days alive) of three rotifers species. For *B. fernandoi*, light pollution significantly affected fecundity (*P*=0.0006; Fig. 2a). Individuals that received light pollution at blue and green spectra exhibited reduced reproduction compared to the treatment without light pollution (μ_*blue*_=3.68±0.536, μ_*green*_=3.76±0.560, μ_*dark*_=7.52±0.988; *P*_*blue-dark*_=0.0037, *P*_*green-dark*_=0.0062). Female offspring production was reduced by 51% and 50% under the blue and green spectra light pollution, respectively. Fecundity per day was also significantly affected by light pollution (*P*<0.001; Fig. 2b). Individuals exposed to white light pollution and without light pollution had the highest fecundity per day (μ_*white*_=0.770±0.078, μ_*dark*_=0.704±0.067) and their fecundity per day was significantly different from that of individuals exposed to blue and green light pollution (μ_*blue*_=0.387±0.041, μ_*green*_=0.448±0.049; *P*_*white-blue*_=0.0001, *P*_*white-green*_=0.0037, *P*_*dark-blue*_=0.0005, *P*_*dark-green*_=0.0191). Despite differences in fecundity, survival was unaffected by light pollution (Fig. 2c).

**Figure 2.**
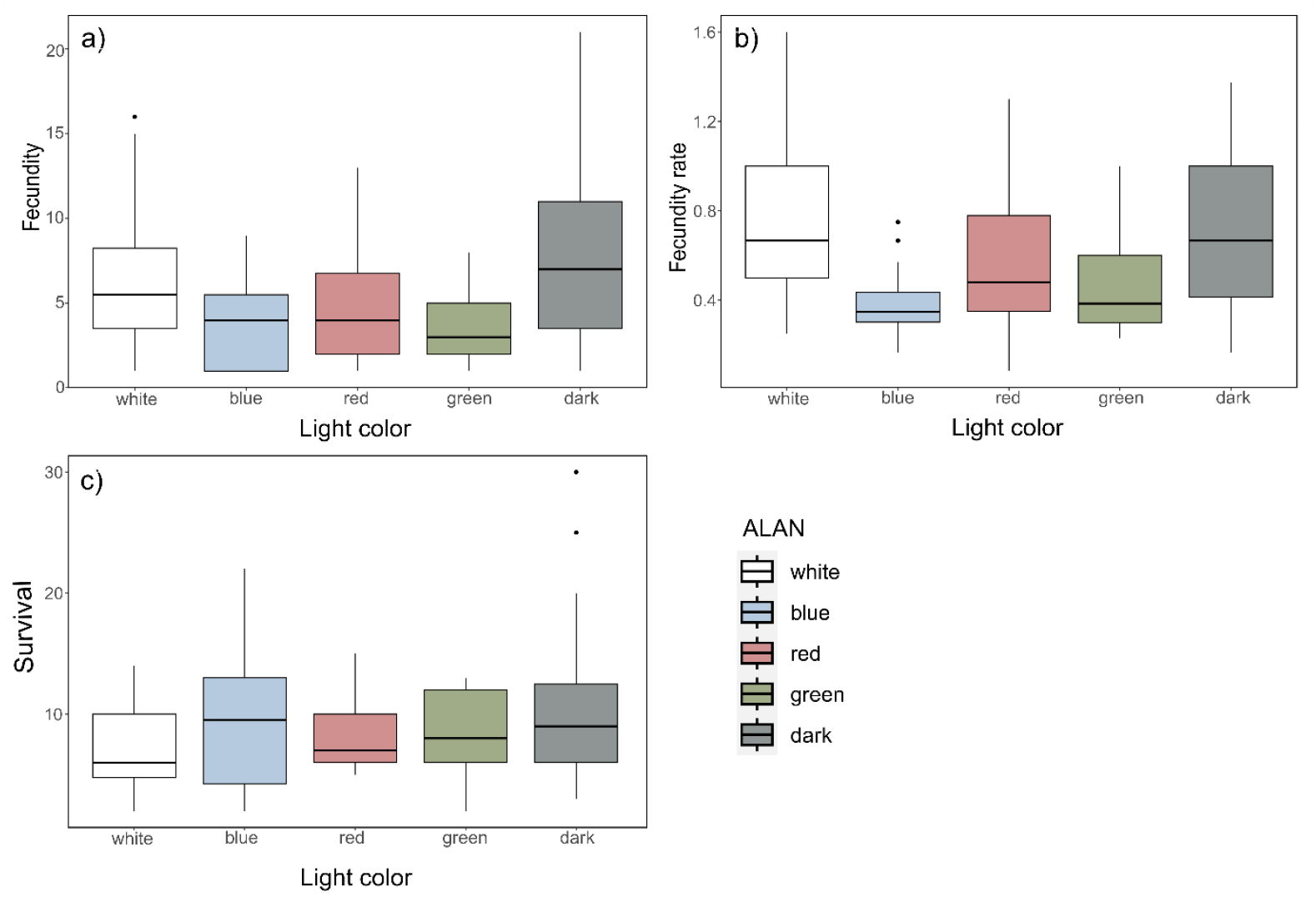
*Brachionus fernandoi* responses to different ALAN spectra. (a) Fecundity of asexual females. (b) Fecundity rate, i.e., fecundity per survival days of asexual females. (c) Survival in days of asexual females. Boxes represent the interquartile range (IQR), and the horizontal line is the median. Whiskers extend up to 1.5 × IQR, points beyond are outliers.

Unlike *B. fernandoi*, in *B. calyciflorus* s.s. both light pollution and rotifer type had a significant effect on survival (*P*_*light*_ <0.0001, *P*_*rotifer_type*_<0.0001; Fig. 3d). The highest survival (in days) was observed in animals exposed to green light pollution, while the lowest survival occurred under white light pollution (μ_*green*_=11.82±1.136, μ_*white*_=5.81±0.568, μ_*dark*_=6.85±0.561, μ_*red*_=6.91±0.606, μ_*blue*_=8.53±0.725; *P*_*green-white*_=0.0001, *P*_*green-dark*_=0.0013, *P*_*green-red*_= 0.0018, *P*_*white-blue*_=0.0225). Sexual animals had a shorter lifespan than asexual ones (μ_*sexual*_=6.51±0.367, μ_*asexual*_=9.45±0.600; P<0.0001). Light pollution also affected the total number of male broods (*P*<0.0001; Fig 3c). More precisely, white light pollution had the lowest number of male broods (μ=2.78±0.318), yet it was significantly different only from green light (μ=5.80±0.630; *P*_*white-green*_=0.0002). Exposure to green light pollution increased the total number of male broods when compared to red light and the treatment without light pollution (*P*_*green-red*_=0.0001, *P*_*green-dark*_=0.0032). The fecundity and fecundity rate of amictic females were unaffected by ALAN (Fig. 3a and 3b).

**Figure 3.**
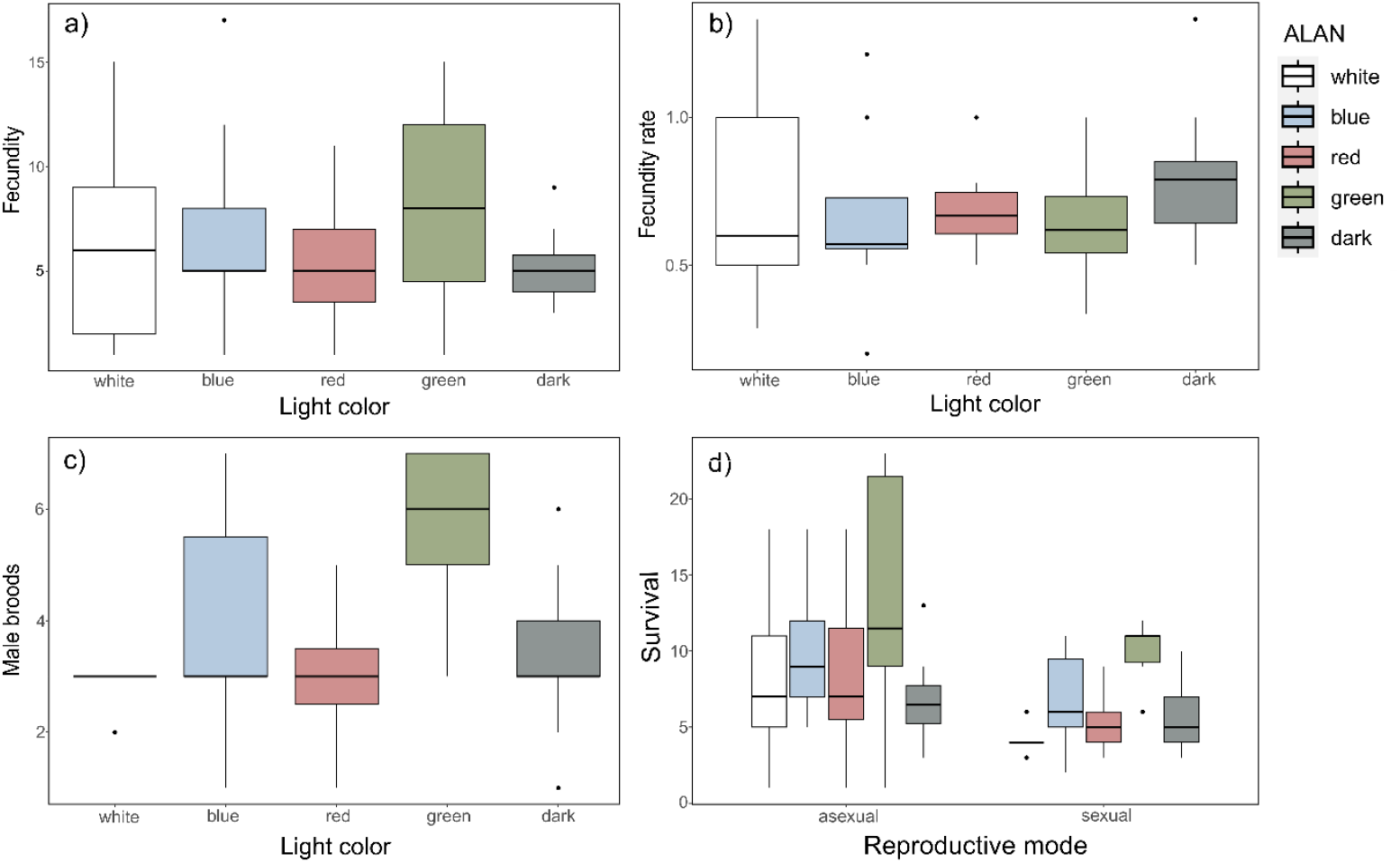
*Brachionus calyciflorus s*.*s*. responses to different ALAN spectra: (a) Fecundity of asexual females. b) Fecundity rate, i.e., fecundity per survival days of asexual females. (c) Production of male broods in sexual females. d) Survival of sexual and asexual females in days. The boxes represent the interquartile range (IQR), horizontal line is the median. Whiskers extend up to 1.5 × IQR, points beyond are outliers.

Similar to *B. fernandoi*, in *B. rubens* light pollution significantly impacted fecundity per day (*P*=0.0174; Fig 4b). Specifically, fecundity per day increased under white light pollution (μ=1.465±0.146) in comparison to blue (μ=0.834±0.103; *P*_*white-blue*_=0.0070), green (μ=0.985±0.061; *P*_*white-green*_=0.0285) and red (μ=0.965±0.097; *P*_*white-red*_=0.0448) light pollution treatments. However, it was not significantly different from the treatment without light pollution (μ=1.026±0.067; *P*_*white-dark*_=0.0611). Although fecundity rate showed significant differences, overall fecundity did not (Fig. 4a). Unlike *B. calyciflorus* s.s., in *B. rubens*, survival was unaffected by light pollution (Fig. 4d). However, asexual females survived longer than sexual ones (μ_*asexual*_=10.95±0.720, μ_*sexual*_=8.23±0.647; *P*=0.0057). Light pollution also affected the total number of male broods (*P*=0.0005; Fig. 4c). More precisely, white light pollution (μ=2.67±0.391) significantly reduced the total number of male broods when compared to blue (μ=4.90±0.429; *P*_*white-blue*_=0.0037), red (μ=4.67±0.391; *P*_*white-red*_=0.0072), green (μ=6.00±0.783; *P*_*white-green*_=0.0042) and no light pollution (μ=4.71±0.512; *P*_*white-dark*_=0.0230) treatments.

**Figure 4.**
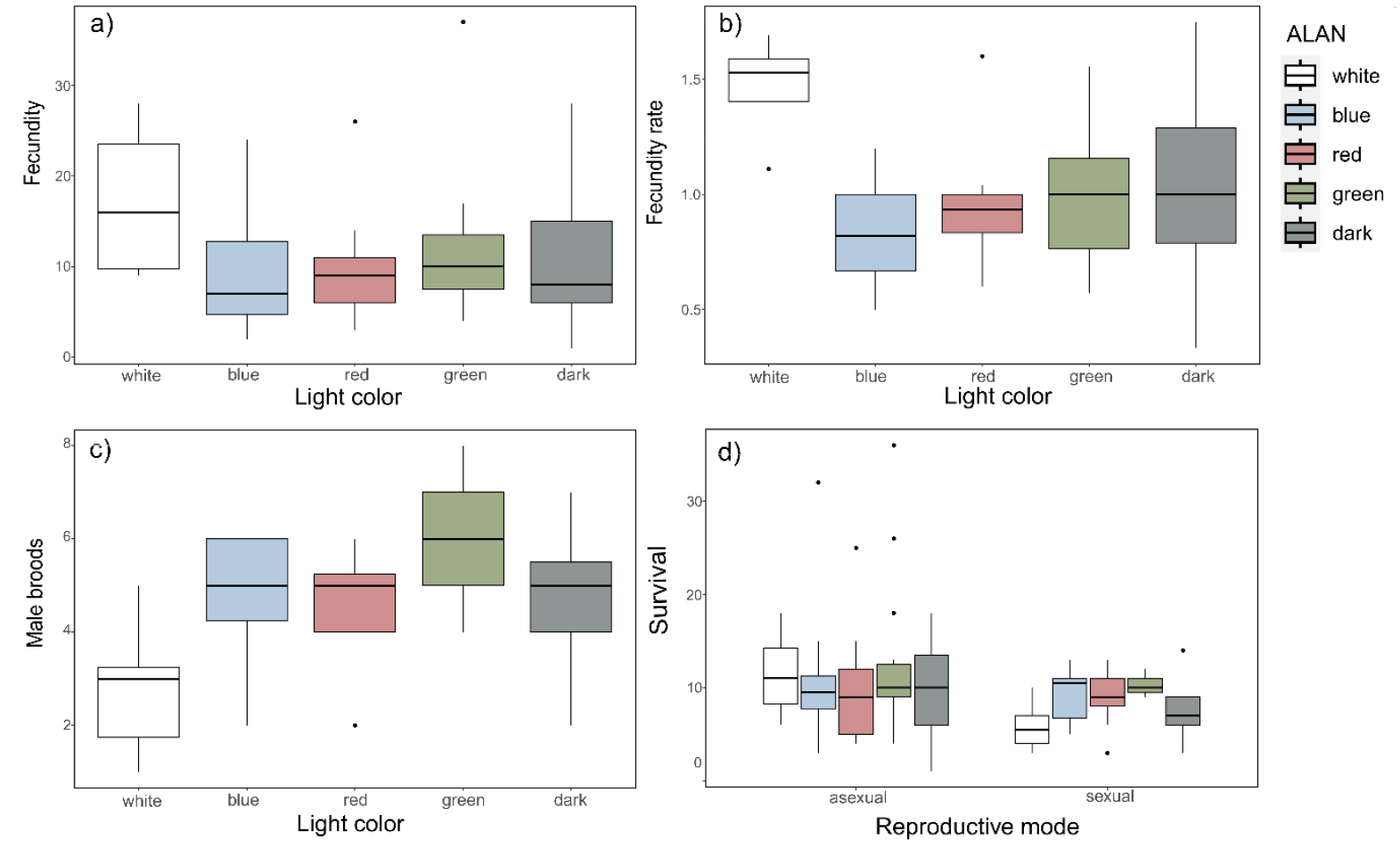
*Brachionus rubens* responses to different ALAN spectra: (a) Fecundity of asexual females. (b) Fecundity rate, i.e., fecundity per survival days of asexual females. (c) Production of male broods in sexual females. d) Survival of sexual and asexual females in days. The boxes represent the interquartile range (IQR), horizontal line is the median. Whiskers extend up to 1.5 × IQR, points beyond are outliers.

### Experiment 2: Effects of light on the attachment behavior of B. rubens

We used ANOVA to analyze the effect of different light spectra on the attachment behavior of *B. rubens*, and found a significant effect of light (F_3,155_=4.204, *P*=0.0068). Specifically, more rotifers attached to *Daphnia* under blue and dark light than to *Daphnia* exposed to daylight (control) (μ_*daylight*_=8.55±0.618, μ_*blue*_=12.55±0.742, μ_*dark*_=11.2±0.687; *P*_*daylight-blue*_=0.0082, *P*_*daylight-dark*_=0.0252; Figure 5).

**Figure 5.**
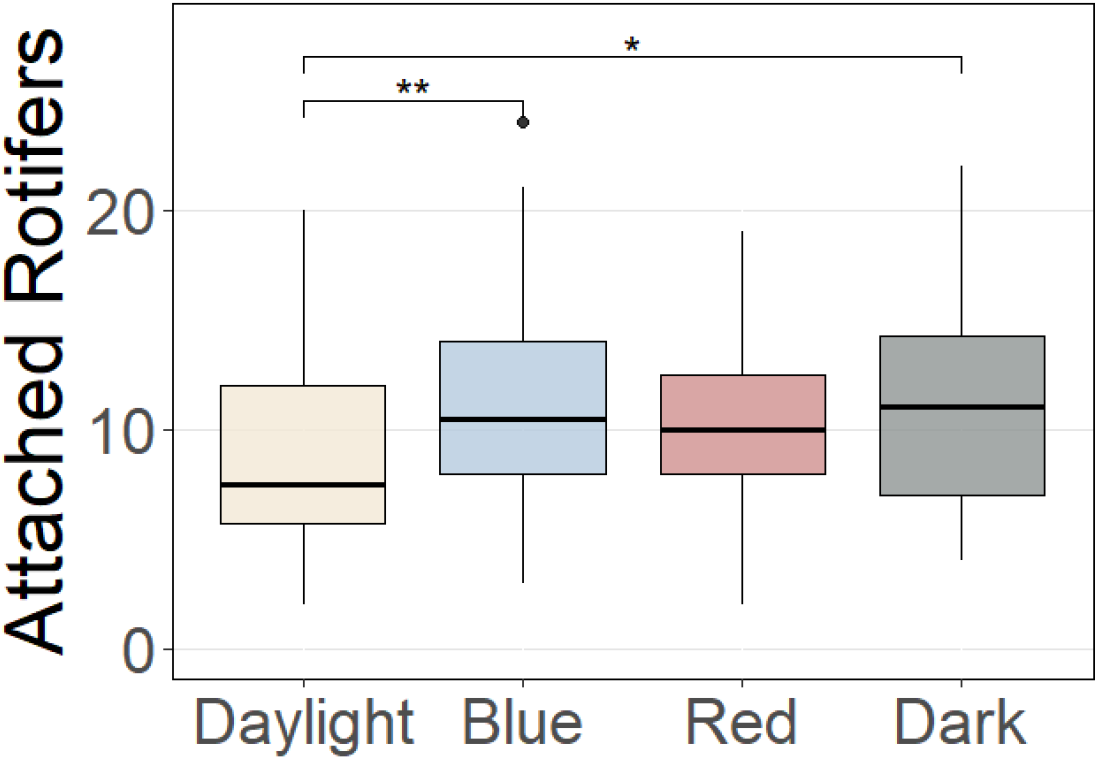
Effect of artificial light on the attachment behavior of *Brachionus rubens*. Asterisks indicate the level of significance (*p-value<0.05, **p-value<0.01). The boxes represent the interquartile range (IQR), horizontal line is the median. Whiskers extend up to 1.5 × IQR, points beyond are outliers.

## Discussion

### Effects of light pollution on life history traits of rotifer species

ALAN aka light pollution is increasingly recognized as a major contributor to global environmental change (Seymoure et al., 2023), with significant effects on aquatic ecosystems and aquatic invertebrates in particular (Ganguly & Candolin, 2023; Grubisic, 2018; Sullivan et al., 2019). In our study, ALAN influenced life history traits differently across three *Brachionus* species. Specifically, green and blue light were the major driver of changes in life history traits, whereas behavioral alterations were more pronounced under white light.

Many biological activities are orchestrated by internal clocks, such as circadian rhythms, which are synchronized by natural light cycles (Gaston et al., 2017). Disruptions to these cycles, such as altered vertical migrations of zooplankton under artificial light (Cohen & Forward, 2009), can have profound effects on the organism’s behavior and consequently its fitness and survival. In numerous invertebrate taxa, ALAN has been shown to disturb circadian rhythms (Duarte et al., 2019; Torres et al., 2020). In rotifers, previous research indicates that light influences their circadian cycles (Sawada & Enesco, 1984; Jiménez-Contreras et al., 2014) and even their survival (Wang et al., 2024). Given that the internal clocks of rotifers regulate food intake and reproduction, disruptions to these rhythms can impair their ability to cope with food stress and trigger inaccurate reproductive cues, ultimately affecting their survival (Bradshaw & Holzapfel, 2007; Yu et al., 2025).

Our findings demonstrate that green light extended the lifespan and male brood production of *B. calyciflorus* s.s., and stand in contrast with studies of *B. plicatilis* where exposure to green ALAN reduced their lifespan (Wang et al., 2024). This discrepancy may be explained by differences in how these species perceive and respond to light via their photoreceptors. For instance, *B. plicatilis* exhibits positive phototaxis towards green light (Kim, Sawada, Rhee, et al., 2014), *i*.*e*., continuous exposure to green ALAN enhances its movement. Increased movement, in turn, raises energetic demands that could trade-off with survival. A study of *B. calyciflorus* sensu lato (s.l.) showed positive phototaxis within the 500-650 nm range, which includes green light (Cornillac et al., 1983). According to the energetic theory, one would expect a similar decline in the lifespan of *B. calyciflorus s*.*s*. Though with recent species delimitation, *B calyciflorus* s.l. comprise four species with distinct ecological differences (*e*.*g*., in temperature tolerance and nutrient requirements; Paraskevopoulou et al., 2018; W. Zhang et al., 2019). Therefore, it is unclear which species responses were observed by Cornillac and colleagues, given that our findings that green ALAN decreased reproduction in *B. fernandoi* corroborate with findings in *B. plicatilis* (Wang et al., 2024).

In *B. fernandoi*, light pollution at the blue spectrum also reduced reproduction. Although no other study has directly linked blue ALAN to reduced reproduction in rotifers, Kim, Sawada, and Hagiwara (2014) reported a decrease in population growth under low-intensity blue ALAN, thereby suggesting that reduced reproduction is likely a key factor, because population growth is driven largely by reproductive output. Typically, reproduction and survival are expected to trade-off against one another, with increased investment in one resulting in a decrease in the other (Sun et al., 2017; Takeshita, 2024). However, we did not observe these trade-offs, neither in *B calyciflorus* nor in *B. fernandoi*. This disruption of the usual survival–reproduction trade-off under green and blue ALAN may indicate that light pollution imposes a stress that alters the normal allocation of resources. The observed variation among different *Brachionus* species underscores the need for further research to measure phototactic responses and energetic costs within the *B. calyciflorus* species complex. Such studies would help clarify how each species responds to ALAN and the underlying physiological and ecological mechanisms driving these responses.

Survival and fecundity are highly dependent on food availability (Schälicke et al., 2019). Some rotifer species, such as *Brachionus urceolaris* and *Brachionus plicatilis*, exhibit twice as high feeding rates during the day compared to the night (Arndt & Heerkloss, 1989; Hammer et al., 2001). Thus, under constant light regime, disruption of the natural photoperiod may lead to nutritional stress, whether by energetic cost or food limitation. In a study by Yu et al. (2025), the adverse effects caused by extreme light regimes to *Brachionus plicatilis*, such as constant light or complete darkness, were mitigated by higher food availability, while more pronounced effects were observed under restricted feeding conditions. Put differently, sufficient food supply increased tolerance to light stress. In our study animals were fed *ad libitum*, ensuring that food limitation did not influence our results. However, the energetic costs associated with increased filtration may still provide an explanation for our findings. The final phenotypic trait we assessed was the number of male broods, which we used as a proxy for male production. Male production decreased during daylight ALAN in *B. rubens*, but increased under green ALAN in *B. calyciflorus* s.s. Although males do not directly contribute to population growth and they are often considered less critical to overall population fitness (W. Li et al., 2022), their production can have important ecological implications, as in many rotifer species it is closely linked to environmental conditions which usually trigger sexual reproduction (Gilbert, 2003; Snell, 2017). A shift towards reduced male production might signal a move towards predominantly parthenogenetic reproduction. While asexual reproduction can support rapid population growth under stable conditions, it can also reduce genetic diversity in the long term, potentially compromising the population’s ability to adapt to future environmental changes. Altered sex ratios could further impact mating dynamics and the frequency of sexual reproduction, thereby affecting the population’s resilience to environmental stressors over time (Schröder & Gilbert, 2004).

### Effects of light on the attachment behavior of B. rubens

Behavioral changes can be induced by exposure to different light spectra, directly impacting an animal’s ability to survive. In our study we focused on a key behavioral trait, the attachment behavior of *B. rubens*, which confers several advantages to rotifers, including refuge from predators or interference by larger daphniids, reduced metabolic costs associated with swimming, and the ability to exploit host metabolic byproducts (Gilbert, 2014; 2019; Iyer & Rao, 1993; 1995) This attachment experiment showed an increase in attachment behavior in all lights tested compared to daylight. Here treatments were applied during daytime hours, thus, rotifers exposed during daytime hours were experiencing a regular daylight cycle, unlike in experiment 1. One initial hypothesis was that disrupted circadian rhythms are responsible for this behavior (Botte et al., 2023). Light can affect both the circadian cycles (Sawada & Enesco, 1984; Jiménez-Contreras et al., 2014) and survival (Wang et al., 2024) in rotifers, thus, changes in light conditions during daytime may disrupt the daily rhythm and circadian clock of rotifers, potentially triggering attachment behavior as a stress response.

Rotifers also show strong phototaxis behavior. For example, *B. plicatilis* exhibits strong positive phototaxis under blue (peaks at 470 nm), green (525 nm) and white (460 and 570 nm) light (Kim et al., 2018). Their photoreceptors are most efficient in absorbing wavelengths between 450 and 550 nm (Kim, Sawada, Hagiwara, 2014; Kim, Sawada, Rhee, et al., 2014), making them highly sensitive to blue and green light. Under blue light, these receptors can become saturated. Saturation reduces sensitivity and diminishes the positive phototactic response (Kim et al., 2018). Thus, a reduction in phototaxis should increase the likelihood of attachment, as rotifers would be less inclined to swim continuously. Despite this being possible in the case of blue light, it does not explain the increase in attachment behavior during the dark treatment.

Despite being clone specific, *Daphnia* hosts have a tendency to swim less during the night (Dodson, 1997). Less swimming activity may facilitate rotifer attachment, since less energy is required to stay attached. Exposure to light increases the number of “hops”, *i*.*e*., a strong power stroke upwards, followed by a short bout of sinking (Sabet et al., 2019). Since these hops require vigorous antennae movement, they could physically dislodge the epibiont *B. rubens*, which could explain why they attach less during daytime compared to nighttime.

## Conclusions

Regardless of the proximate mechanism, alterations in attachment behavior carry both ecological and evolutionary implications. In natural environments, decreased attachment could expose rotifers to higher predation risk and force them to invest more energy in continuous swimming, thereby creating potential trade-offs in energy allocation. Although our current experiments did not reveal immediate fitness deficits, chronic ALAN exposure might eventually affect growth or reproductive output through transgenerational effects. Over evolutionary time, persistent ALAN might exert selective pressure on the photoreceptive systems of rotifers, favoring individuals with photoreceptors that are less prone to saturation or that exhibit modified phototactic thresholds, potentially leading to evolutionary shifts in sensory physiology and attachment behavior. It is also important to note that ALAN affected the three *Brachionus* species differently, suggesting that no single mechanism is at work. Nevertheless, it is clear that ALAN alters species interactions, potentially triggering cascades of ecological effects, feedback responses and time lags that can lead to complex shifts within communities. Understanding these dynamics is crucial, as even subtle disruptions in light regimes may have far-reaching consequences for aquatic ecosystem structure and function.

## Acknowledgments

We thank Claus Peter Stelzer for providing us with resting eggs from Egelsee, Austria that were used to establish the *B. calyciflorus* s.s. clone. We thank Imad Rabah for her contribution in data collection and Noga Kronfeld-Schor for lending us a spectrometer to measure and adjust the desirable wavelength frequencies for each light treatment.

## Funding Source Acknowledgments

S.P. was supported by a Fellowship provided by the Steinhard Natural History Museum and a joint Fellowship provided by Tel Aviv University, Israel and the University of Potsdam, Germany.

## Author Contributions

I.R collected the data. E.B. and S.P. analyzed, interpreted the data and wrote the manuscript with input from I.R and F. B-A. S.P led data collection. F. B-A and I.R. conceptualized the study.

