## Supplementary material for "Light pollution affects the behavior and life history traits of aquatic invertebrates"

### A. Natural Day light

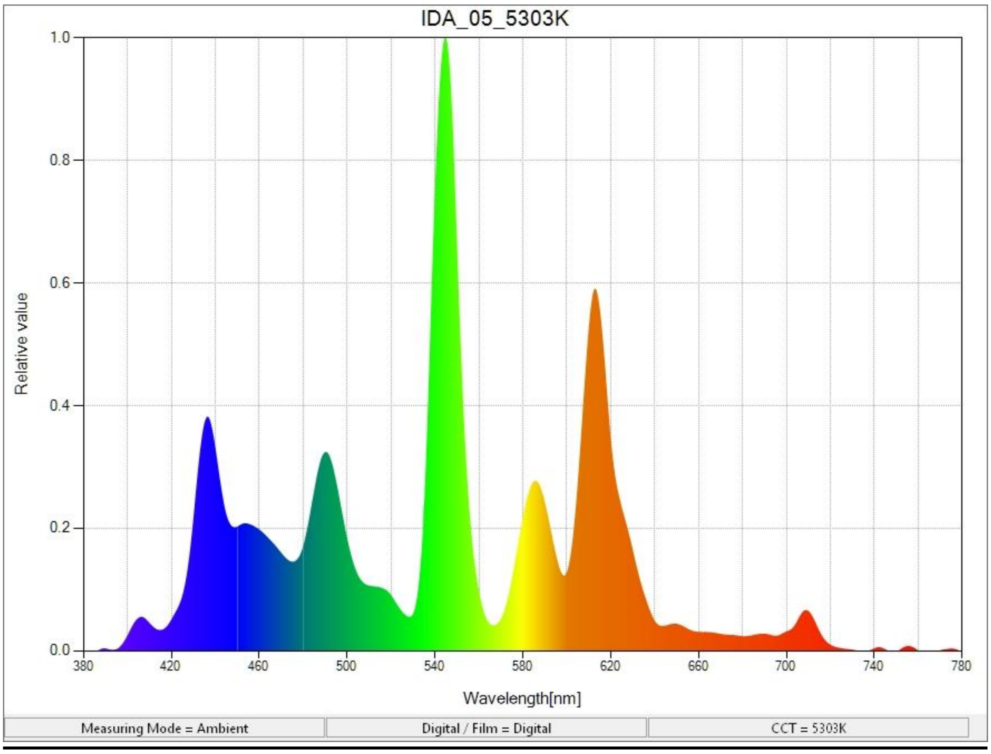

B. Treatment white light (Life history experiment)

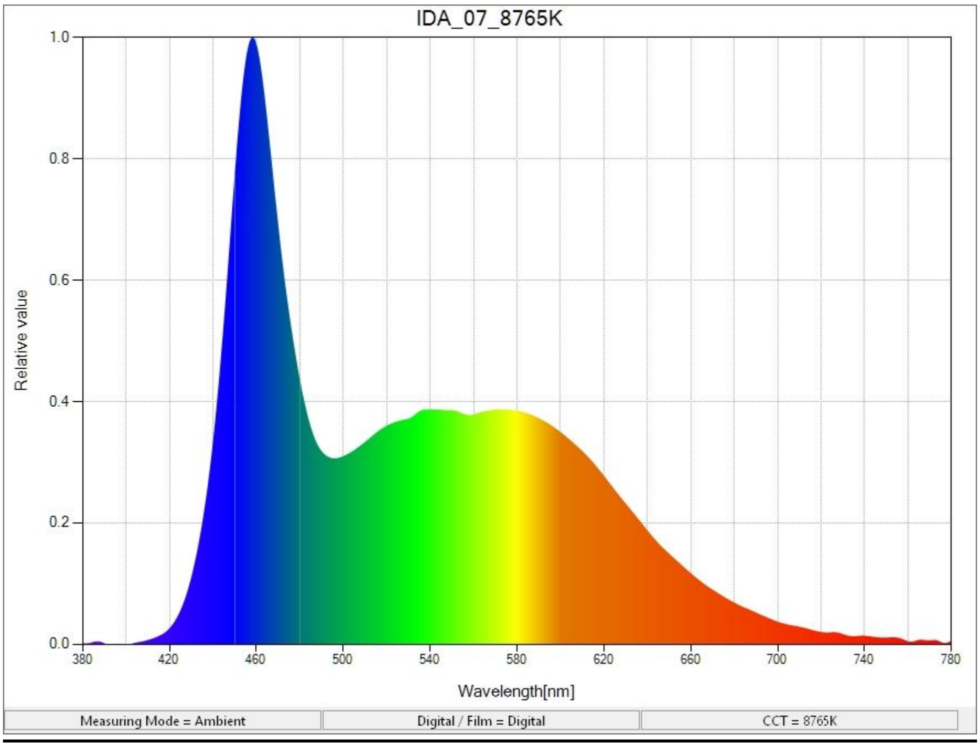


C. Red

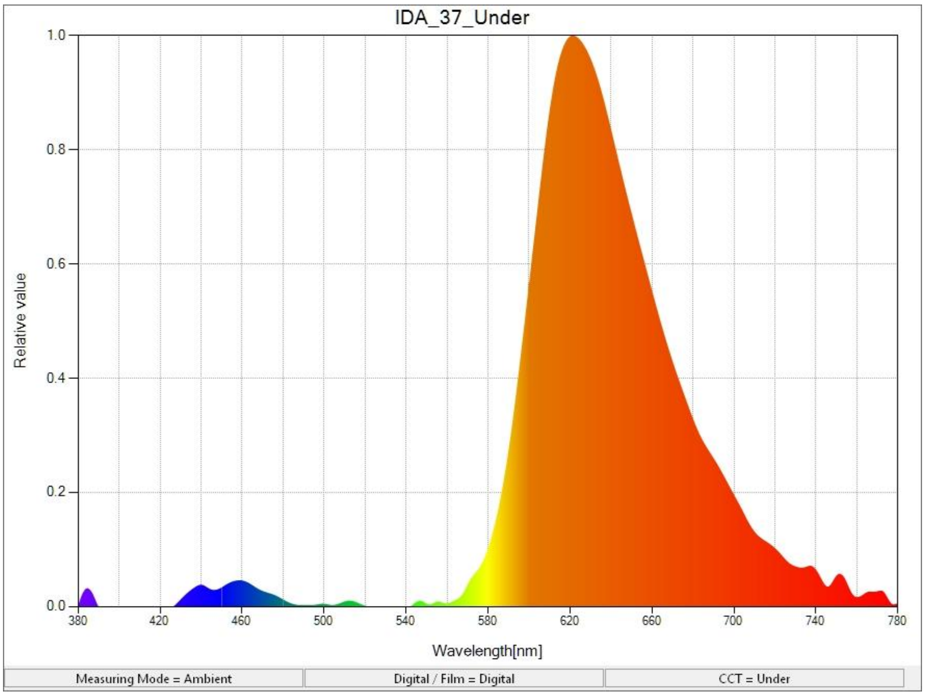

D. Yellow

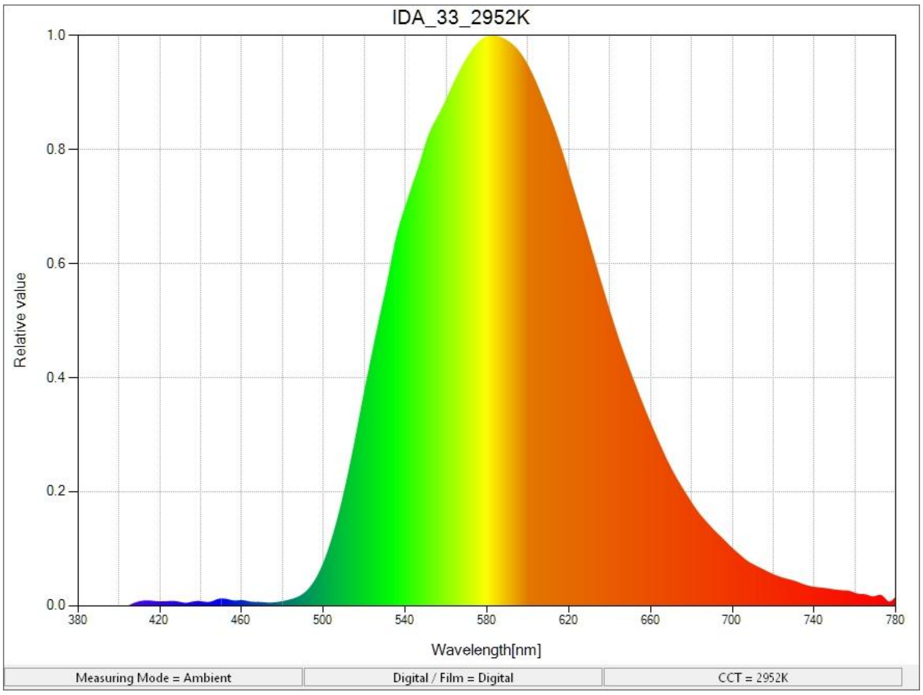


E. Green

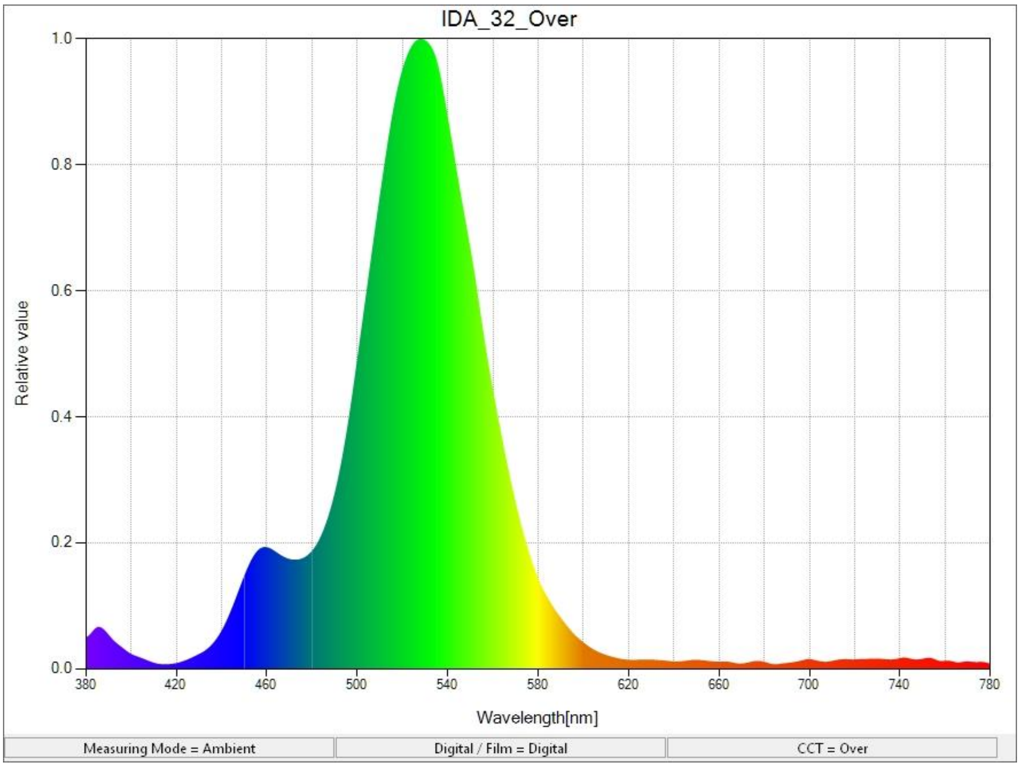

F. Blue

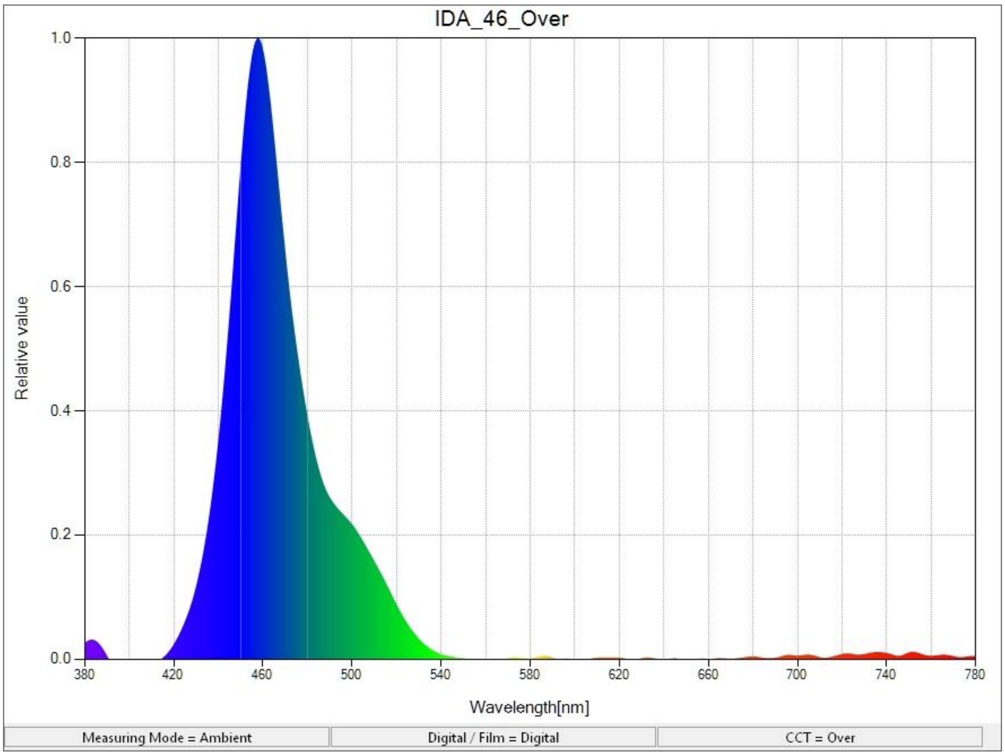


**Figure S1.** Spectral distribution plot that shows the intensity of light at different wavelengths A) natural day light, B) white light treatment, C) red light treatment, D) yellow light treatment, E) green light treatment, and F) blue light treatment**.**
